# Ability to predict irregular periods of food deprivation improves body-weight regulation and reduces weight gain in food-insecure starlings

**DOI:** 10.1101/2025.04.02.646744

**Authors:** Charlotte Parker, Ryan Nolan, Clare P. Andrews, Melissa Bateson

**Author notes:** Author for correspondence, Address: Biosciences Institute, University of Newcastle, Henry Wellcome Building, Framlington Place, Newcastle upon Tyne, NE2 4HH, UK.

## Abstract

Food insecurity is associated with higher body weight in humans and other species, but the causal effect of unpredictable food availability on weight gain is unknown. We measured food intake and weight in starlings (*Sturnus vulgaris*) exposed to repeated irregular periods of food deprivation. We manipulated the predictability of deprivation between subjects with a 1-hour visual cue that either reliably preceded deprivation (Predictable), or was uncorrelated with deprivation (Unpredictable). During the cue, Predictable birds reduced their food intake and spent less time inactive, indicating that they had learnt the contingency. Despite these responses, they lost less weight during subsequent deprivation. They also ate less and gained less weight when food was returned. Birds with the largest behavioural response to the cue had the lowest overall variance in body weight. Consistent with the insurance hypothesis, food intake and body weight increased over time in both groups and body weight was higher in the Unpredictable group. Our results suggest that when food deprivation was predictable, birds were less reliant on stored fat and instead used conditioned hypometabolism to mitigate the effects of food deprivation. We discuss the implications of our findings for the differential health impacts of food insecurity and intermittent fasting.

## 2. Introduction

Food insecurity—defined as limited and unpredictable access to nutritionally adequate food—is associated with obesity and poor health in humans (1,2). In contrast, intermittent fasting, which, like food insecurity can involve entire days without eating, is associated with weight loss and improved health (3–5). We hypothesise that the explanation for this apparent paradox lies in the distinction between starvation and fasting (6): whereas starvation is driven by extrinsic constraints, that are often unpredictable, fasting is internally driven and predictable. In the current paper, we test the role of predictability in determining the impact of irregular food deprivation on food consumption and body weight in European starlings (*Sturnus vulgaris*), an animal model of food insecurity-induced weight gain (7–9).

The insurance hypothesis suggests that food insecurity-induced weight gain is the result of evolved mechanisms that trigger additional fat storage as a buffer against starvation when cues of food scarcity are detected (10,11). Optimality models show that fat stores should be higher when access to food is more unpredictable (12–14). Supporting these models, experiments in several species, including starlings, show that irregular periods of unpredictable food deprivation cause fat storage and weight gain (8,15). However, these experiments confound irregular food deprivation with unpredictability of food deprivation and thus shed no light on the role played by unpredictability *per se* in food insecurity-induced weight gain (16).

There are strong reasons to believe that predictable schedules of food availability should improve regulation of body weight and metabolic health. Expectation of food can trigger preparatory physiological responses that facilitate homeostasis (17). For example, the anticipatory rise in insulin triggered by the taste of sugar in the mouth, or even the sight of food, functions to rapidly stabilise blood glucose following food intake (18). Animals can also learn to mitigate the impact of food deprivation by increasing their food intake prior to removal of food (19–22) and by rapidly initiating hypometabolism in response to food deprivation before any energy deficit occurs (23,24). Thus, animals possess flexible learning mechanisms that deliver adaptive physiological and behavioural responses to anticipated changes in food availability. We hypothesise that the disruption of these mechanisms when schedules of food intake are unpredictable could explain the food insecurity-intermittent fasting paradox. Individuals able to predict deprivation should be able to mitigate its effects with short-term adjustments to food intake or energy expenditure, whereas individuals unable to predict deprivation must permanently carry additional fat stores as insurance, at a cost to long-term health (11).

Here we present an experiment on starlings with either cued (Predictable) or uncued (Unpredictable) food deprivation, designed to test the role of unpredictability in food insecurity-induced weight gain. We randomly allocated birds to one of two groups. In the Predictable group, the food bowl changed colour, from black to green, for one hour immediately prior to irregular periods of food deprivation. In the Unpredictable group, the same cue was equally likely to occur, either one hour prior to deprivation, or for a matched hour on a day when no deprivation occurred, and was thus uncorrelated with deprivation. The only difference between the groups was therefore the information available to the birds: both groups experienced an irregular schedule of periods of food deprivation interspersed with *ad libitum* access to food, but for the Predictable group, the timing of deprivation was predictable, if they learnt the contingency associated with the cue.

The design was based on a previous experiment in which starlings failed to learn the cue (16). We made three modifications to this experiment to increase the likelihood that birds in the present Predictable group would learn the cue. First, to limit the potential for birds to use time of day as a cue, the period of deprivation started at one of eight possible randomly chosen times and to avoid social cuing the start time was different for each bird on a given day. Second, to increase the salience of the cue, we used the colour of the food bowl. We predicted that this cue would be more likely to be associated with food deprivation than the ambient light cue used previously, because it was entirely novel to the birds and spatially adjacent to food. Third, to further increase both the salience of the cue and give the birds more time to respond to it, we doubled the duration of the cue to 1 hour. The primary dependent variables we recorded were food intake, dawn body mass and the feeding and perching behaviour of the birds in response to the cue.

We predicted that birds in the Predictable group would learn to respond to the cue by immediately increasing their food intake as a short-term strategy for buffering upcoming deprivation. Consequently, while both groups would show an increase in food intake following deprivation, this rebound would be larger in the Unpredictable group, because they would have a larger energy deficit to make up. Second, we predicted that both groups would gain weight over the course of the experiment as insurance against the starvation risks associated with irregular food deprivation, but that weight gain would be higher in the Unpredictable group, due to their greater degree of uncertainty and hence requirement for greater insurance. These predictions were the same as those made previously and were therefore preregistered in van Berkel *et al*. (16).

## 3. Materials and Methods

### Design

The experiment was a between-subjects design with birds randomly allocated to one of two treatments: Predictable and Unpredictable food deprivation. Birds in both treatments experienced 16 repeated 5-hour periods of food deprivation spread irregularly over 32 days and at all other times, food was available *ad libitum*. In the Predictable treatment (n = 8 birds), a cue was presented that perfectly predicted upcoming food deprivation. In the Unpredictable treatment (n = 8), the same cue was presented an equal number of times, but the cue was uncorrelated with upcoming food deprivation (Figure 1).

**Figure 1.**
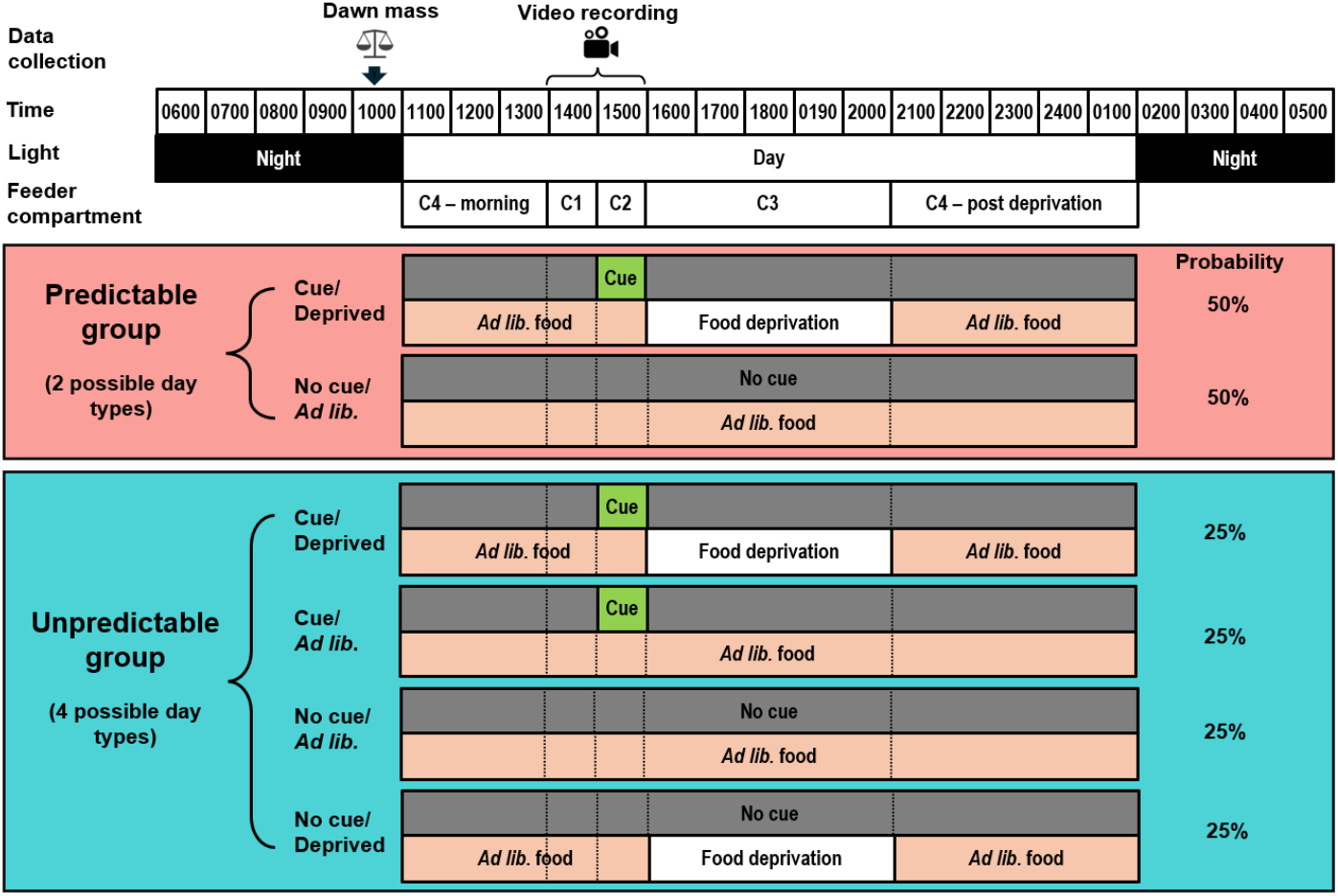
Experimental design. Over the 32-day experiment, birds were exposed to a 5-hour period of food deprivation on 50% of days (deprived days). Food deprivation and replacement was accomplished using programmable, rotating feeders with four compartments (C1-C4) assigned to the different periods of the day (C4 covered the period before C1 and after C3). In order that birds could not use time of day to anticipate deprivation, there were eight, equally probable daily schedules, with the deprivation period (C3) starting at any hour between 13:00 and 20:00 (in the example schedule shown, deprivation starts at 16:00). During this deprivation period, on deprived days birds received no food, while on *ad libitum* days food was *ad libitum*. The deprivation period was always directly preceded by a 1-hour cue period (C2) and the cue period was preceded by a 1-hour baseline period (C1). When the cue was present in C2, the compartment was coloured green. The vertical dotted lines indicate the times at which the feeder rotated to reveal the next compartment. Thus, two types of day were possible in the Predictable group and four in the Unpredictable group. Birds received the different schedules and day types in a semi-random order.

### Animals

The subjects were 16 adult European starlings (6 female, 10 male) originally taken from wild nests on day 5 post-hatching and hand-reared in the laboratory as part of a previous study that involved manipulating food intake and begging effort during development (25). This prior manipulation was not part of the current study and was fully counterbalanced along with natal nest between the two treatment groups in the current experiment. The birds were 4 years old at the time of the current study. Immediately prior to the current experiment, the birds were group-housed in an indoor aviary supplied with *ad libitum* food and water (see details below). After the completion of the current experiment, the birds were retained in the laboratory for further studies.

### Experimental room and husbandry

For the experiment, replicates of 8 birds were transferred to an experimental room. Ambient temperature and humidity matched the conditions in the holding aviary (∼18°C, ∼40% humidity). The light schedule was 15 hours light: 9 hours dark with incremental brightening/dimming over 30-minutes at either end of the day to simulate dawn/dusk (birds were clock-shifted prior to the start of the experiment to facilitate pre-dawn weighing and husbandry; full daylight began at 1100). Birds were housed in individual cages (74 × 45 × 45cm), arranged so that they had auditory and visual contact. Each cage contained two wooden perches, two water drinkers (supplemented with vitamin supplement; BSP vitamin drops, Vetark Professional), floor papers, and an automated feeder (see below). A nutritionally complete diet, Special Diets Services Poultry Starter (HPS), henceforth ‘food’, was available *ad libitum* except during periods of food deprivation (see below), supplemented with four live mealworms each day. This diet was also fed for three days prior to catching birds from the holding aviary to reduce the habituation necessary when the birds were initially transferred to experimental cages.

Daily husbandry in the experimental room was commenced just prior to dawn onset in order to minimise disturbance of the birds’ foraging time and ensure that measurements of the previous day’s food consumption were accurate (birds do not eat in the dark). Automated feeders were removed from the cages. Remaining food was weighed and feeders were cleaned and reset in accordance with the individual feeding schedule for each bird.

Husbandry was carried out and each cage was supplied with a water bath for the dawn period. Five minutes before the start of full daylight, baths were replaced with the feeders, birds were given their mealworms and video cameras were set for the day. From full daylight onwards, there was no further entry into the experimental room until the following morning.

### Manipulation of food deprivation and cue presentation

Each cage was equipped with an automated feeder (Andrew James 4 Meal Programmable Automatic Pet Feeder) that provided all of the bird’s food for the duration of the experiment. The feeder had four removeable compartments (C1-C4) that could each be filled with 20-g food (a quantity that was more than a bird could eat during the maximum available period for a compartment) and programmed to become sequentially accessible at any hour in the 24-hour period (Table 1). The compartments rotated such that when a new compartment became accessible, the previously available compartment became inaccessible, allowing accurate measurement of consumption during the period each compartment was accessible.

**Table 1.** Details of the four compartments in the automated feeders.

| Compartment | Description | Start time range* | Duration | Food amount |
| --- | --- | --- | --- | --- |
| C1 | **Baseline period** | 1100-1800 | 1 hour | 20 g every day |
| C2 | **Cue period** (cue only present in C2 on 50% of days). On cue days C2 was coloured green. | 1200-1900 | 1 hour | 20 g every day |
| C3 | **Deprivation period** (deprivation only occurred during C3 on 50% of days) | 1300-2000 | 5 hours | 20 g on the 50% of days that were *ad lib*. No food on 50% of days that were deprived. |
| C4 | Remainder of the day (split between before the baseline period and after the deprivation period on most days) | Dawn until C1 then after C3 until dusk. | 8 hours light (+ subsequent dark period when birds don't eat) | 20 g every day |

\*There were 8 possible start times for each compartment (always on the hour). Each of the 8 schedules was experienced 4 times by each bird.

The food compartments were made of black plastic and were deep enough to allow the colour to be clearly visible above the level of the food. A novel colour cue was created by covering the inner surface of a compartment with green PVC tape. On cue days only, this cue was placed in C2.

### Procedure

#### Allocation of birds to experimental treatments

The experiment ran in two sequential replicates, with each replicate comprising four birds in each treatment group. At the start of each replicate, eight birds were caught from the holding aviary and transferred to individual cages in the experimental room. In both replicates, the number of males and females was unequal, meaning that while two males and two females were allocated to the Predictable group, three males and one female were allocated to the Unpredictable group. Cages were arranged on two double-level shelving units and birds were allocated to cages such that equal numbers from each natal nest and treatment group were on the top and bottom shelves. Within the constraints of counterbalancing, allocation of birds to treatments was random.

Following transfer to cages, the birds were allowed to habituate to their new environment and the automated feeders for 1 week. During this period, *ad libitum* food was continuously available from the automated feeders (20 g in each of the four compartments). The compartments rotated on random schedules to habituate the birds to the apparatus. The experimental phase started once daily food consumption had stabilised for all individuals.

### Experimental Phase

The experimental phase lasted for 32 days and comprised 16 *ad libitum* days and 16 deprived days randomly distributed with the constraint that there were never more than two consecutive deprived days. On *ad libitum* days, birds always had food available. On deprived days, birds had a 5-hour period of total food deprivation starting at any hour between 1300 hours and 2000 hours (yielding 8 possible schedules). Birds experienced each deprivation schedule twice, with the order of schedules randomised for each bird.

On 16 days the cue was present in C2 and on the other 16 days there was no cue present. In the Predictable group, the cue was always presented in the 1-hour cue period immediately prior to the deprivation period on deprived days only; the cue was never presented on *ad libitum* days. In the Unpredictable group, the cue was presented in the 1-hour cue period, immediately prior to deprivation on 8 deprived days (a randomly chosen day of each schedule) and in a matched period on 8 *ad libitum* days (see Figure 1).

Thus, the Predictable group was exposed to an irregular but perfectly predictable schedule of food deprivation. In contrast, the Unpredictable group were exposed to the same irregular schedule of deprivation and the same number of cue presentations, but the cue provided no information about upcoming deprivation.

### Dependent variables

#### Dawn body mass

Birds were caught and weighed just before dawn, when the gut was empty, every three or four days throughout the experimental phase, with the first mass on day 1 and the last on day 33. We obtained a total of 10 weight measurements for each bird.

#### Food intake

Food intake from each compartment in the feeder was measured daily during pre-dawn husbandry. We calculated total daily food intake as the sum of the amount consumed from the four compartments.

#### Feeding bouts and perching inactive

We filmed the birds on 4 deprived and 4 *ad libitum* days (with pairs of days matched for feeder schedule) spread over the second half of the experiment (days 17-32). We used Behavioural Observation Research Interactive Software (26) to manually score two behavioural variables during the baseline (C1) and cue (C2) periods: total number of feeding bouts and proportion of time spent perching inactive. A feeding bout was a behavioural event recorded by continuous observation, defined as occurring when a bird that was perched on the edge of the feeder moved its head fully below the rim of the food bowl. Perching inactive was a behavioural state, defined as occurring when a bird had both feet on the perch and the wings folded; perching was recorded by time sampling at 30-second intervals (27). We computed both variables for the baseline and cue periods for each bird on each available day. Due to camera failure and data loss, video from 7.63 ± 1.09 (mean ± SD) days per bird was available for the eating analysis and 7.13 ± 1.15 days per bird for the perching analysis.

### Statistical Analysis

Data were analysed in R (version 4.2.3). Data files and R scripts are available on the Open Science Framework (http://osf.io/vyp4r). We used linear mixed models (LMM) fitted using the package ‘lme4’. Assumptions of normality and homogeneity of variance were confirmed by visual inspection of model residuals.

Bird (a factor with 16 levels) was included as a random effect in all models that involved repeated measurements of individual birds. Although natal nest was also a potential source of non-independence in the data, natal nest was excluded from the random effects in the final models presented, because it frequently explained zero variance leading to singularities in the model fitting. Natal nest could not be a confound, because natal nest was fully counterbalanced in the experimental design.

Since male starlings are heavier on average than females and therefore also eat more, we included sex (factor: Male/Female) in all models to control for the unbalanced number of males and females in the Predictable and Unpredictable groups. The main independent variables included in each model are detailed in the Results section. These included: experimental treatment (factor: Predictable/Unpredictable); day of experiment (integer: 1-33); recency of deprivation (defined as the number of *ad libitum* days since the last deprived day; factor: 0, 1 or 2+ where 0 corresponds to deprivation the previous day in analyses of dawn body mass, but deprivation on the current day in analyses of food intake); and total daily food intake (continuous). Continuous predictor variables were centred to facilitate interpretation of parameter estimates in models including interactions.

In confirmatory analyses to test preregistered predictions, we used type III ANOVA computed with Satterthwaite’s method for statistical inference and adopted the standard criterion for significance of p < 0.05. In exploratory analyses, given the number of potential variables and interactions, we used a model selection approach implemented in the R package ‘MuMin’ as an alternative approach to inference (28). By convention, we retained all models within 2 AICc units of the top model in our best-models subset. Where more than one model was retained, we then used model averaging to obtain parameter estimates for the supported independent variables. To express the degree of support for a variable we summed the total AIC weight received by models including that term; 100% support indicates that the variable was supported by all the models in the best-models subset.

## 4. Results

### The Predictable group reduced their food intake in response to the cue

We measured food intake in four consecutive periods each day: baseline period (C1; 1 hour), cue period (C2; 1 hour), deprivation period (C3; 5 hours during which birds received no food on deprived days and unlimited food on *ad libitum* days; Figure 1) and post-deprivation period (C4; 8 hours; Figure 2A). To test our predictions regarding the acute effects of deprivation on food intake, we fitted LMMs to food intake in each period, with treatment (Predictable/Unpredictable), day type (*ad libitum*/deprived) and their interaction as independent variables (see Table S1 for ANOVA results). In the cue period, there was a significant interaction between treatment and day type (F_1,493_ = 72.90, p < 0.001), but contrary to our prediction, this was driven by Predictable birds eating significantly less on deprived days (post-hoc pairwise comparison on deprived days: t_13_ = 2.57, p = 0.023). In the post-deprivation period, as predicted, there was a significant main effect of day type (F_1,493_=178.98, p < 0.001), due to both groups eating more following deprivation on days on which they had been deprived. There was a marginally non-significant interaction between treatment and day type (F_1,493_=3.51, p = 0.062), suggesting that as predicted, this rebound was larger in the Unpredictable group (post-hoc pairwise comparison on deprived days: t_13_ = 2.61, p = 0.022).

**Figure 2.**
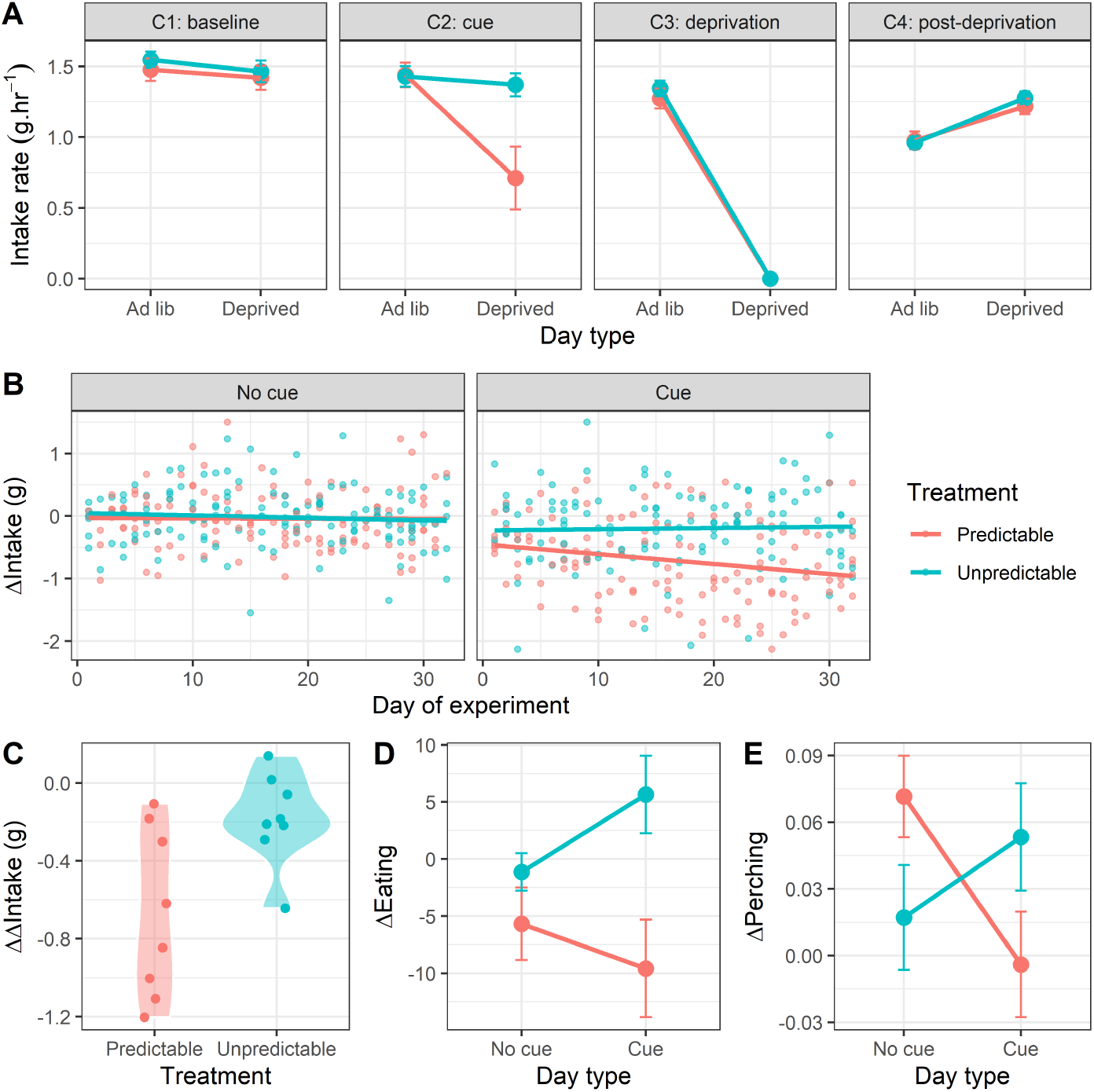
The Predictable group learnt the cue and reduced their food intake during it. (A) Food intake rate (g.hr^-1^) during consecutive periods of the day corresponding to the four compartments of the automated feeders on *ad libitum* days (when no deprivation occurred) and deprived days (when birds were deprived for 5 hours during C3). For C4, we used intake data from the subset of days on which the majority of C4 (≥ 5 hours) fell after the deprivation period. Points are mean intake rates for 8 birds in each treatment ± 1 SE. (B) Change in rate of food intake between the baseline period and cue period (ΔIntake). The facets show plots from days with no cue and days when the cue was presented. Points are daily values for individual birds. Negative values indicate that the rate of intake declined during the cue period relative to the baseline period. (C) Difference in Intake (ΔΔIntake) between days with and without the cue. Points represent the mean values for individual birds across all 32 days of the experiment. Negative values indicate a greater decline in intake between the baseline and cue period on days when the cue was present compared to days when it was not presented. (D) Change in the total number of feeding bouts between the baseline and cue period (ΔEating). Negative values indicate that number of feeding bouts declined during the cue period relative to the baseline period. Points are mean values for 8 birds in each treatment ± 1 SE. (E) Change in the proportion of time samples on which birds were perching inactive between the baseline and cue period (Perching). Negative values indicate that proportion of time perching declined during the cue period relative to the baseline period. Points are mean values for 8 birds in each treatment ± 1 SE.

Our design was predicated on the birds in the Predictable group learning that a change in the colour of the food bowl predicted a period of deprivation commencing an hour later. The results presented above strongly suggest that the Predictable group responded to the cue, albeit in the opposite direction to our prediction. To test this more directly and control for the decline in rate of intake across the day evident in Figure 2A, we computed a measure of behaviour change in response to the presentation of the cue by calculating the difference in intake between the baseline period and the cue period (ΔIntake = Intake_C2_ - Intake_C1_), whereby a positive value indicates that food intake increased during the cue period. To test our predictions regarding the effects of cue presence on ΔIntake we fitted an LMM with treatment, cue presence (cue/no cue), day of experiment (1-32) and all interactions as independent variables (Table S2). There was a significant 3-way interaction between treatment, cue presence and day of experiment (F_1,489_ = 4.43, p = 0.036): in line with the result presented above, the birds in the Predictable group responded to the cue by reducing their rate of intake during the cue and this effect increased with time indicative of learning (Figure 2B). A simple slopes analysis of the data from the cue days showed that ΔIntake was stable over time in the Unpredictable group (LMM: β = 0.00, t = 0.34, p = 0.736), but declined significantly over time in the Predictable group (LMM: β = -0.02, t = -3.49, p < 0.001).

To derive a single metric of learning for each bird, we calculated the difference in mean ΔIntake between cue and no cue days (ΔΔIntake = ΔIntake_cue_ - ΔIntake_no cue_) using the data from all 32 days. More negative values indicate a greater reduction in intake in response to the cue on days when the cue was present and hence constitute stronger evidence of learning; birds that learned sooner and/or had a larger response to the cue should have a more negative ΔΔIntake. As expected, given that learning could only occur in the Predictable group, ΔΔIntake was significantly more negative in the Predictable group (two-sample t-test: t_11_ = -2.82, p = 0.017; Figure 2C).

To provide additional evidence of learning, we scored the behaviour of the birds from video recorded on 8 days in the second half of the experiment. We measured the total number of feeding bouts and proportion of time perching inactive during the baseline and cue periods for each bird on each available day. From these data, we calculated ΔEating and ΔPerching, where positive values indicate, respectively, an increase in the number of feeding bouts and the proportion of time perching during the cue period relative to the baseline period. To test the effect of cue presence on ΔEating and ΔPerching, we fitted LMMs with treatment, cue presence and their interaction as independent variables (Tables S3 and S4; day was not included in these models, because we only had video data from the second half of the experiment). In both models, there was a significant interaction between treatment and cue presence (ΔEating: F_1,107_ = 5.05, p = 0.026; ΔPerching: F_1,102_ = 9.04, p = 0.003): Predictable birds responded to the cue by reducing the number of bouts of feeding and by reducing the proportion of time spent perching inactive (Figure 2D, E). These results provide corroborating evidence that the Predictable group learnt the cue and that they responded to it by reducing food consumption. The perching result suggests that the Predictable birds did not respond to the cue by becoming more inactive.

### The Predictable group had more stable body weights and were lighter overall

On day 1 of the experiment there was no significant difference in dawn body mass between birds in the Predictable and Unpredictable groups (two-sample t-test: t_13_ = -0.60, p = 0.558). To test our predictions regarding the cumulative effects of irregular deprivation on dawn mass, we fitted an LMM with treatment, day of experiment and their interaction as independent variables (Table S5). As predicted, there was a significant linear effect of day on dawn mass (F_1,142_ = 8.32, p = 0.005), resulting in a mean increase of 1.43 g (1.88%) over the 32 days of the experiment (Figure 3A). Contrary to our predictions, there was no evidence that treatment affected the rate of mass increase (F_1,142_ = 0.418, p = 0.519).

**Figure 3.**
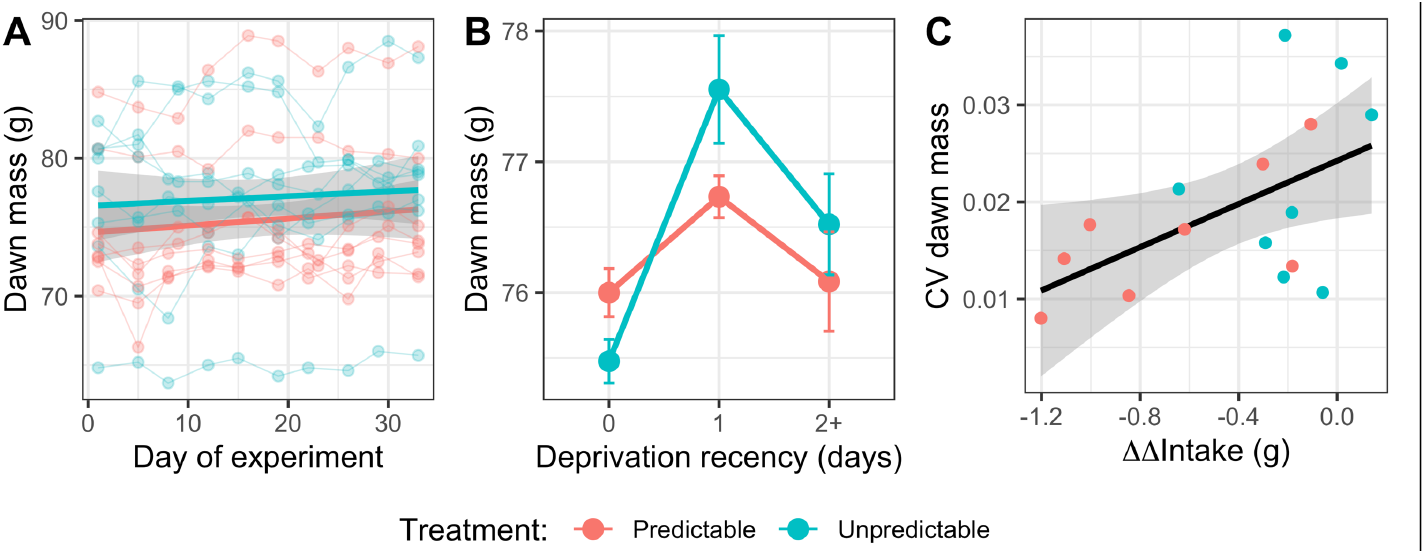
The Predictable group had more stable body weight and were lighter overall. (A) Cumulative effect of irregular food deprivation: body weight by day of experiment. Points are individual measurements and thin lines connect points from a bird. Linear regression lines for the two treatment groups are shown with 95% confidence intervals. (B) Acute effect of deprivation: body weight by deprivation recency. Points are means ± 1 SE for 8 birds in each treatment. Data were within-subject centred and then expressed relative to the grand mean to facilitate visualisation of within-subject effects. (C) Coefficient of variation (CV) of dawn mass for each bird by the behavioural response to the cue (ΔΔIntake; a metric of learning). Points are the overall values for each of the 16 birds. The linear regression is shown with 95% confidence intervals.

To explore the acute effects of deprivation on dawn mass, due to the number of potential interactions and the absence of specific hypotheses, we used information theoretic model selection for inference. We specified a maximal LMM with effects of treatment, day of experiment, deprivation recency (last deprivation 0, 1 or 2+ days ago) and all possible interactions. This yielded a total of 256 alternative models (with sex also included). Three top models emerged with a combined AIC weight of 60% (Table S6). To understand the effects of the independent variables supported, we computed model-averaged parameter estimates from these three models (Table S7). The model selection showed strong support for a non-linear effect of deprivation recency (summed AIC weight = 100%): birds that had not been deprived for two or more days were heavier than birds with 0 *ad libitum* days since last deprivation and lighter than birds last deprived 1 day previously (Figure 3B). Thus, birds had lost weight on the day immediately following food deprivation, regained additional weight on the next day after returning to *ad libitum* food, then partially lost this additional weight over subsequent days with *ad libitum* food. Moreover, there was support for an interaction between treatment and deprivation recency (summed AIC weight = 66%): Predictable birds lost less weight on days they were food deprived and regained less weight following 1 day of *ad libitum* recovery. Replicating the results above, there was support for a positive linear effect of day of experiment (summed AIC weight = 63%). In addition, as originally predicted, there was strong support for a main effect of treatment (summed AIC weight = 100%), with Unpredictable birds being heavier overall when acute effects of deprivation recency were controlled for in the model.

Finally, we explored whether an individual bird’s variance in body weight in response to deprivation was explained by how much it reduced its rate of intake in response to the cue. To characterise variance in dawn mass, for each bird we computed the coefficient of variation (CV) in dawn mass across all 32 days. Birds with more negative values of ΔΔIntake (i.e. birds with a larger overall reduction in rate of food intake in response to the cue) had significantly lower CVs in dawn mass (GLM: β ± se = 0.01 ± 0.00, F1,13 = 5.62, p = 0.033; Figure 3C).

### Daily intake increased over the experiment, but the Predictable group ate less overall

On day 1 of the experiment there was no significant difference in total daily food intake between birds in the Predictable and Unpredictable groups (two-sample t-test: t_13_ = -0.42, p = 0.680). To explore the effects of deprivation on total daily food intake we used model selection. We specified a maximal LMM with effects of treatment, day of experiment, deprivation recency and all possible interactions (sex was also included). Of the 256 alternatives, three top models emerged with a combined AIC weight of 78% (Tables S8 and S9). The model selection showed strong support for a positive linear effect of day of experiment (summed AIC weight = 100%): birds in both groups increased their daily food intake over the 32 days of the manipulation by 2.04 g equating to a 17.6% increase in daily intake (Figure 4A). There was also strong support for a non-linear effect of deprivation recency (summed AIC weight = 100%): birds that had not been deprived for two or more days ate more than birds deprived that day, but less than birds deprived one day ago. Moreover, the top model showed support for an interaction between treatment and deprivation recency (summed AIC weight = 46%): Predictable birds ate less on days they were food deprived and less on the following *ad libitum* day (Figure 4B).

**Figure 4.**
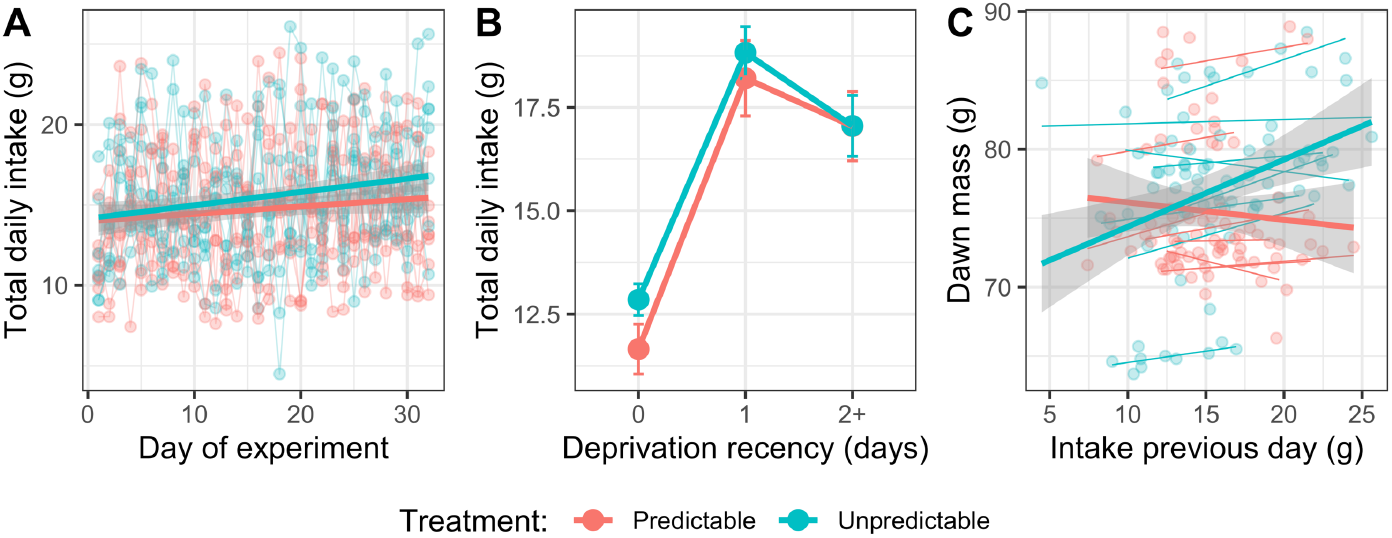
Daily food intake increased over the experiment, but the Predictable group ate less overall. (A) Total daily food intake by day of experiment. Points represent individual measurements and thin lines connect points from a bird. Overall fitted lines for the two treatment groups are shown. (B) Total daily food intake (mean ± SE) by deprivation recency. (C) Dawn mass by food intake the previous day. Panels A and C show linear regression lines for the two treatment groups with 95% confidence intervals.

To explore the contribution made by food intake to variation in dawn body mass, we fitted a LMM to dawn mass with total intake the previous day, treatment, and their two-way interaction as independent variables (Table S10). Eating more food was associated with a significantly higher body mass the following morning (F_1,102_ = 18.07, p < 0.001). The main effect of treatment and the interaction between treatment and intake were not significant, although a plot of the data suggested that the association between intake and dawn mass was weaker in the Predictable group, consistent with the results reported above (Figure 4C).

## 5. Discussion

We tested how the predictability of irregular periods of food deprivation affected food intake and body weight in European starlings. Our aim was to determine whether the same schedule of irregular food deprivation caused greater weight gain when deprivation was unpredictable to the birds, compared with when they could predict it 1 hour in advance. We found that birds able to predict deprivation had more stable weights, ate less and were lighter overall. Our results suggested that the mechanism limiting weight loss during deprivation in the Predictable group was likely to be a conditioned reduction in resting energy expenditure. Below we discuss our findings and explore their potential relevance for understanding how humans and other animals respond to food deprivation experienced during food insecurity and intermittent fasting.

### Acute effects of predictability on behaviour and body mass

The Predictable group changed their rate of food intake in response to the cue and this change in behaviour increased in magnitude over the experiment, consistent with the birds learning the contingency associated with cue (Figure 2). Thus, our manipulation succeeded in creating groups of animals for whom schedules of food deprivation differed only in predictability; confounds present in previous designs, including exposure to the cue and irregularity and severity of deprivation, were held constant between the groups (8,29,30) (see also discussion in (16)).

Contrary to our prediction of increased food intake in anticipation of deprivation, the Predictable birds showed a decline over the experiment in their rate of food intake during the 1-hour cue period (Figure 2B). Given that Pavlovian conditioning is a learning mechanism designed to deliver adaptive anticipation of fitness-relevant stimuli (31), this response appears maladaptive. However, the unconditional response to starvation in many species including birds and humans is energy-saving hypometabolism, characterised by reduced resting metabolic rate and physical activity (6). Furthermore, animals can learn to initiate hypometabolism before experiencing an energy deficit resulting in reduced weight loss during subsequent food deprivation (23,24). It is therefore possible that stopping eating and initiating hypometabolism during the cue period is the response that delivered the smallest overall energy deficit in our experiment. This could be the case if hypometabolism takes time to initiate and the subsequent energy savings during deprivation more than compensate for the additional food that could be consumed during the cue period. The fact that the Predictable birds lost less weight on days they were food deprived (Figure 3B), despite eating less than the Unpredictable birds (Figure 4B), suggests that their response to the cue resulted in net energy savings. Flying vertebrates have smaller intestines and shorter gut retention times compared to non-flying animals (32), which may favour hypometabolism over anticipatory eating as a short-term strategy for coping with imminent food deprivation in passerine birds due to constraints on their ability to binge in preparation for deprivation (but see (19,33) for evidence of anticipatory eating in finch species).

The Predictable birds also reduced the proportion of time they spent perching inactive during the cue period, suggesting that their energy savings were not the result of reduced physical activity. Diurnal hypothermia in the absence of reduced physical activity has been described in quail exposed to food deprivation, demonstrating that birds can save energy by hypothermia whilst remaining active (23). Further work is required to establish whether conditioned hypothermia was responsible for the improved resistance to food deprivation shown by the Predictable group in the current experiment. In summary, we have shown that the information provided by the cue allowed the Predictable birds to anticipate food deprivation and reduce their resting energy expenditure, thereby limiting the weight loss they sustained during deprivation.

As predicted, birds in both groups compensated for food deprivation by eating more, both immediately after deprivation and in total on the following *ad libitum* day. There was also support for the predicted treatment difference in the size of this rebound intake, with Unpredictable birds consuming more following deprivation (Figure 2A and C). This result makes sense, because since the Predictable birds lost less weight during deprivation, the energy deficit to be made up afterwards was smaller. In line with this result, our exploratory analyses showed that the Predictable birds also gained less weight on the first *ad libitum* day following deprivation.

Thus, Predictable birds had more stable body weight over a cycle of food deprivation and recovery and the evidence suggests that this was driven by a reduction in energy expenditure, most likely lower resting metabolic rate. Moreover, individual birds that showed the strongest behavioural evidence for learning the predictability of deprivation also had the least variable body masses over the experiment, suggesting that predictability improved the homeostatic regulation of body weight during a fast. In contrast, the Unpredictable birds, who could not anticipate deprivation, showed a more exaggerated, cyclic, “yo-yo” pattern of weight loss and regain. Weight cycling has been demonstrated to cause increased fat deposition and a rise in biomarkers of metabolic dysfunction in mice (34).

### Cumulative impact of unpredictability on behaviour and body mass

In accordance with predictions from the insurance hypothesis, both groups of birds increased their dawn mass over the experiment and the Unpredictable group were on average heavier when we controlled statistically for deprivation recency. This result replicates extensive previous work in starlings and other species including mice, showing that unpredictable food deprivation causes rapid increase in body fat and/or weight (7,8,15,35). This response makes adaptive sense, since both groups experienced unpredictable periods of food deprivation in the early stages of the experiment before the Predictable group had learned the cue, but the uncertainty, and hence the requirement for additional fat as insurance, was higher overall in the Unpredictable group.

Although there was statistical support for the Unpredictable group being heavier overall, we did not find the expected interaction between treatment group and time: there was no evidence that the rate of weight gain was higher in the Unpredictable group over the 32 days of the experiment (Figure 3A). It is possible that this interaction was not detected because, to reduce additional stress, we only caught and weighed the birds every 3-4 days and individual weight measurements were often impacted by recent deprivation. In contrast, food intake was measured daily, and we saw an interaction between treatment and time on total daily food intake, with total daily intake increasing at a higher rate in the Unpredictable group (Figure 4A). Given our finding that total daily food intake predicted body weight the following morning (Figure 4C), we would have expected to see the treatment difference in food intake mirrored in the body weights, and the fact that we did not is probably a limitation of our measurement protocol. In future studies we suggest measuring weight daily using a less stressful method (e.g. 9) and adding 2-3 *ad libitum* days for all birds at the end of the experiment to ensure that final weight measurements are not acutely affected by recent deprivation (see 29).

### Implications for food insecurity and intermittent fasting

Food insecurity and intermittent fasting are both characterised by irregular, periods without eating. For example, in humans, reporting periodically not eating for entire days is a criterion for a classification of severe food insecurity (36) and fasting for two non-sequential 24-hour periods in every seven days (the 5:2 diet) is one of the available protocols for intermittent fasting (5). Given this similarity, it seems paradoxical that food insecurity and intermittent fasting are associated with opposite effects on body weight, health and longevity. While we acknowledge that there are likely to be many differences between individuals experiencing food insecurity and those who voluntarily engage in intermittent fasting (e.g., socioeconomic status, diet quality), our current results suggest that the ability to predict when food deprivation will occur could contribute to the different health outcomes. Food insecurity is defined by uncertainty and associated anxiety about future access to food (37), whereas intermittent fasting is within the control of the participant and hence totally predictable. This insight could explain some of the variation in outcomes from both animal and human trials of intermittent fasting. For example, mice exposed to two days a week of fasting (supposedly modelling the 5:2 diet) reported none of the positive effects on body weight, health and longevity that are typically associated with alternate-day fasting (38,39). We speculate that this fasting schedule would be hard for the mice to learn and might have therefore simulated food insecurity as opposed to intermittent fasting. It is an open question whether humans might also find some irregular fasting schedules harder to learn than others (e.g., 5:2 diet versus alternate-day fasting or time restricted eating), impacting their efficacy for weight reduction.

## 6. Conclusions

Our results show that the impact of irregular periods of food deprivation depends on whether the deprivation is predictable in advance. Food deprivation that was unpredictable caused more exaggerated weight cycling and higher overall body weight in starlings. In birds that could predict deprivation, our results suggest that information about upcoming deprivation was sufficient to trigger a reduction in resting metabolic rate that reduced weight loss during deprivation and hence the fat stores birds needed to carry as insurance against starvation.

These findings contribute to our understanding of the biological impacts of food insecurity and support the hypothesis that unpredictable food deprivation disrupts normal homeostatic control of body mass that occurs via conditioned regulation of metabolic rate. We conclude that cognition related to food availability—including potentially unconscious learning of eating schedules—may be critical in explaining the differential health impacts of food insecurity and intermittent fasting.

## Acknowledgments

We thank Daniel Nettle for his support throughout the project and for comments on the manuscript.

## Ethical Statement

The study adhered to ASAB/ABS guidelines for the use of animals in research. Birds were taken from the wild under Natural England permit 20121066 and the research was completed under UK Home Office licence PPL 70/8089 with approval of the Animal Welfare and Ethical Review Body at Newcastle University.

## Data Accessibility

The data, meta-data and R code necessary to generate the analyses and figures in the current paper have been deposited on the Open Science Framework and are publicly available as of the date of submission at: http://osf.io/vyp4r.

## Authors’ Contributions

Conceptualization and Methodology C.P., R.N., C.P.A. and M.B.; Investigation C.P. and C.P.A.; Data curation C.P.; Formal analysis C.P., R.N., C.P.A. and M.B.; Visualization C.P. and M.B.; Supervision C.P.A. and M.B.; Project Administration M.B.; Writing – original draft C.P. & M.B.; Writing – review & editing C.P., R.N., C.P.A. and M.B.

## Supplemental information

**Table S1.** ANOVA results from LMMs testing the effects of deprivation on food consumption.

| Independent variable | Dependent variable |  |  |  |  |  |  |  |
| --- | --- | --- | --- | --- | --- | --- | --- | --- |
|  | Baseline: C1 intake rate (g.hr<sup>-1</sup>) |  | Cue period: C2 intake rate (g.hr<sup>-1</sup>) |  | Deprivation period: C3 intake rate (g.hr<sup>-1</sup>) |  | Post-deprivation period: C4 intake rate (g.hr<sup>-1</sup>)† |  |
|  | F ratio | P value | F ratio | P value | F ratio | P value | F ratio | P value |
| Treatment | F<sub>1,13</sub>=0.01 | 0.941 | F<sub>1,13</sub>=3.13 | 0.100 | F<sub>1,13</sub>=0.11 | 0.742 | F<sub>1,13</sub>=0.02 | 0.902 |
| Day type | F<sub>1,493</sub>=4.56 | **0.033\*** | F<sub>1,493</sub>=100.37 | **<0.001\*\*\*** | F<sub>1,493</sub>=5811.66 | **<0.001\*\*\*** | F<sub>1,493</sub>=178.98 | **<0.001\*\*\*** |
| Treatment* Day type | F<sub>1,493</sub>=0.14 | 0.706 | F<sub>1,493</sub>=72.90 | **<0.001\*\*\*** | F<sub>1,493</sub>=4.29 | **0.039\*** | F<sub>1,493</sub>=3.51 | 0.062 . |
| Sex | F<sub>1,13</sub>=4.35 | 0.057 . | F<sub>1,13</sub>=0.42 | 0.527 | F<sub>1,13</sub>=3.98 | 0.068 . | F<sub>1,13</sub>=2.57 | 0.133 |

\*\*\*p < 0.001, \*p < 0.050, .p < 0.100.
†For this analysis we used consumption data from the subset of days on which the majority of C4 (≥ 5 hours) fell after the deprivation period.

**Table S2.** ANOVA results from LMM testing whether ΔIntake showed evidence for cue learning.

| Independent variable | F ratio and df | P value |
| --- | --- | --- |
| Day of experiment | F<sub>1,489</sub> = 3.41 | 0.065 . |
| Treatment | F<sub>1,13</sub> = 5.82 | **0.031 \*** |
| Cue presence | F<sub>1,489</sub> = 88.90 | **< 0.001 \*\*\*** |
| Sex | F<sub>1,13</sub> = 0.51 | 0.488 |
| Day of experiment * Treatment | F<sub>1,489</sub> = 2.13 | 0.145 |
| Day of experiment * Cue presence | F<sub>1,489</sub> = 0.95 | 0.329 |
| Treatment * Cue presence | F<sub>1,489</sub> = 30.47 | **< 0.001 \*\*\*** |
| Day of experiment * Treatment * Cue presence | F<sub>1,489</sub> = 4.43 | **0.036 \*** |

\*\*\*p < 0.001, \*p < 0.050, .p < 0.100.

**Table S3.**
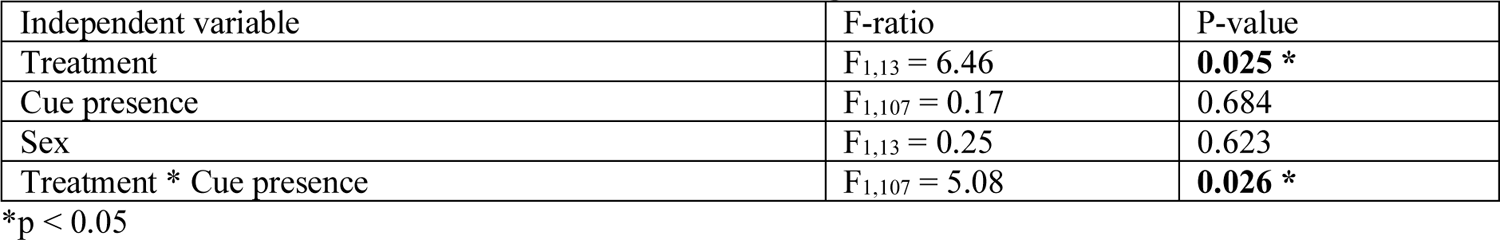
ANOVA results from LMM on ΔEating.

| Independent variable | F-ratio | P-value |
| --- | --- | --- |
| Treatment | F<sub>1,13</sub> = 6.46 | **0.025 \*** |
| Cue presence | F<sub>1,107</sub> = 0.17 | 0.684 |
| Sex | F<sub>1,13</sub> = 0.25 | 0.623 |
| Treatment * Cue presence | F<sub>1,107</sub> = 5.08 | **0.026 \*** |

\*p < 0.05

**Table S4.** ANOVA results from LMM on ΔPerching.

| Independent variable | F-ratio | P-value |
| --- | --- | --- |
| Treatment | F<sub>1,14</sub> = 0.01 | 0.918 |
| Cue presence | F<sub>1,102</sub> = 0.90 | 0.346 |
| Sex | F<sub>1,14</sub> = 0.09 | 0.7702 |
| Treatment * Cue presence | F<sub>1,102</sub> = 9.036 | **0.003 \*\*** |

\*\*p < 0.01

**Table S5.** ANOVA results from LMM on dawn mass.

| Independent variable | F-ratio | P-value |
| --- | --- | --- |
| Day of experiment | F<sub>1,142</sub> = 8.32 | **0.005 \*\*** |
| Treatment | F<sub>1,13</sub> = 0.105 | 0.752 |
| Sex | F<sub>1,13</sub> = 0.86 | 0.372 |
| Day of experiment * Treatment | F<sub>1,142</sub> = 0.418 | 0.519 |

\*\*p < 0.01

**Table S6.**
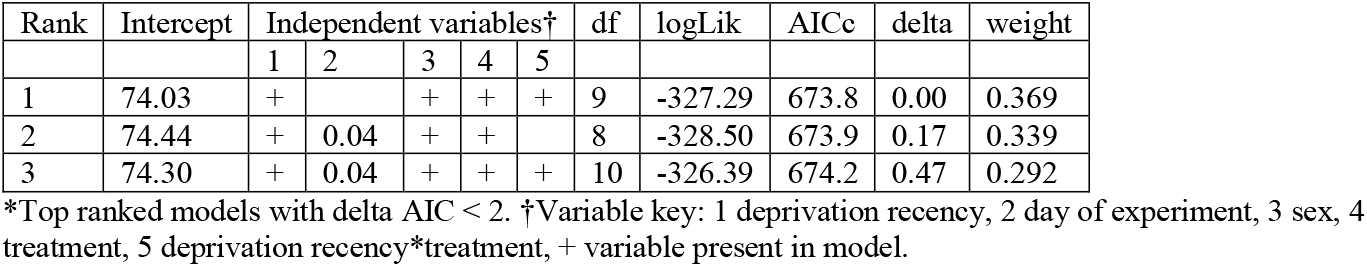
Model selection table for effects on dawn mass.*

**Table S7.** Model-averaged parameter estimates for supported effects on dawn mass.

| Independent variable | Reference category | Estimate† | SE | Summed AIC weights % |
| --- | --- | --- | --- | --- |
| Intercept |  | 74.25 | 2.55 |  |
| Deprivation recency[0] | 2+ | -0.69 | 0.55 | 100 |
| Deprivation recency[1] | 2+ | 0.22 | 0.43 | 100 |
| Sex[male] | female | 2.87 | 3.05 | 100 |
| Treatment[Unpredictable] | Predictable | 1.13 | 2.97 | 100 |
| Deprivation recency[0]*Treatment[Unpredictable] | Predictable | -0.50 | 0.62 | 66 |
| Deprivation recency[1]*Treatment[Unpredictable] | Predictable | 0.31 | 0.58 | 66 |
| Day of experiment |  | 0.03 | 0.02 | 63 |

†Values are the full averages of estimates from the three models listed in Table S8.

**Table S8.** Model selection table for effects on total daily food intake.*

| Rank | Intercept | Independent variables† |  |  |  |  | df | logLik | AICc | delta | weight |
| --- | --- | --- | --- | --- | --- | --- | --- | --- | --- | --- | --- |
|  |  | 1 | 2 | 3 | 4 | 5 |  |  |  |  |  |
| 1 | 16.24 | + | 0.06 | + | + | + | 10 | -1101.70 | 2223.84 | 0.00 | 0.46 |
| 2 | 16.05 | + | 0.06 | + |  |  | 7 | -1105.25 | 2224.72 | 0.88 | 0.30 |
| 3 | 15.92 | + | 0.06 | + | + |  | 8 | -1104.42 | 2225.12 | 1.28 | 0.24 |

\*Top ranked models with delta AIC < 2. †Variable key: 1 deprivation recency, 2 day of experiment, 3 sex, 4 treatment, 5 deprivation recency\*treatment, + variable present in model.

**Table S9.** Model-averaged parameter estimates for supported effects on daily intake.

| Independent variable | Reference category | Estimate† | SE | Summed AIC weights (%) |
| --- | --- | --- | --- | --- |
| Intercept |  | 16.10 | 0.74 |  |
| Deprivation recency[0] | 2+ | -5.11 | 0.39 | 100 |
| Deprivation recency[1] | 2+ | 1.27 | 0.33 | 100 |
| Sex[male] | female | 1.70 | 0.85 | 100 |
| Treatment[Unpredictable] | Predictable | -0.01 | 0.78 | 68 |
| Deprivation recency[0]*Treatment[Unpredictable] | Predictable | 0.45 | 0.59 | 46 |
| Deprivation recency[1]*Treatment[Unpredictable] | Predictable | 0.16 | 0.39 | 46 |
| Day of experiment |  | 0.06 | 0.01 | 100 |

†Values are the full averages of estimates from the three models listed in Table S10.

**Table S10.** ANOVA results from LMM on dawn mass exploring effects of daily food intake.

| Independent variable | F-ratio | P-value |
| --- | --- | --- |
| Treatment | $F_{1,143} = 9.79$ | **<0.002 \*\*** |
| Total intake on previous day | $F_{1,13} = 0.11$ | 0.750 |
| Sex | $F_{1,13} = 0.73$ | 0.409 |
| Total intake on previous day * Treatment | $F_{1,143} = 0.11$ | 0.741 |

\*\*p < 0.01.

